# Development of a Microelectrode Array System for Simultaneous Measurement of Field Potential and Glutamate Release in Brain Slices

**DOI:** 10.1101/2025.03.30.646242

**Authors:** Aiko Hasegawa, Naoki Matsuda, Ikuro Suzuki

**Affiliations:** Department of Electronics, Graduate School of Engineering, Tohoku Institute of Technology, 35-1 Yagiyama Kasumicho, Taihaku-ku, Sendai, Miyagi, 982-8577, Japan

**Author notes:** Corresponding author: Ikuro Suzuki.

**Keywords:** Microelectrode array, Enzyme-modified carbon nanotubes electrode, simultaneous FP/EC measurement system, Glutamate, Brain slice

## Abstract

Disorders of the central nervous system and complex side effects caused by abnormal neurotransmitter release have been widely reported. If neurotransmitter release and field potential (FP) could be simultaneously measured in real time, it would be possible to capture the relationship between changes in neurotransmitter release and alterations in electrical activity. In this study, we developed a novel microelectrode array (MEA) system capable of simultaneously measuring FP and electrochemical (EC) signals. Additionally, we developed an enzyme-modified carbon nanotube (CNT)-MEA capable of detecting glutamate, a major neurotransmitter, with high sensitivity in the range of several nM to several hundred nM. Using the developed enzyme-modified CNT-MEA and the simultaneous FP/EC measurement system, we successfully recorded extracellular potentials and glutamate release from hippocampal brain slices, detecting both an increase in oscillatory activity and temporal changes in glutamate release following caffeine administration. Furthermore, we also achieved the recording of dopamine release from brain slices using this system. We believe that the enzyme-modified CNT-MEA and the simultaneous FP/EC measurement system, which enable the real-time, simultaneous measurement of FP, glutamate release, and dopamine release, will contribute to a deeper understanding of brain circuit mechanisms, pathological brain conditions, and the evaluation of pharmacological compounds.

**Highlights:**

- The MEA system was customized to facilitate real-time simultaneous measurement of field potential (FP) and electrochemical (EC) signals.
- The enzyme-modified CNT-MEA demonstrated the capability to detect glutamate at concentrations ranging from a few nM to several hundred nM.
- Using the enzyme-modified CNT-MEA and the simultaneous FP/EC measurement system, we concurrently recorded FP and glutamate release from hippocampal brain slices, as well as their dynamic changes induced by caffeine administration.
- The developed enzyme-modified CNT-MEA successfully detected dopamine release from brain slices.
- The simultaneous recording of neural activity, glutamate release, and dopamine release achieved in this study is expected to contribute to elucidating the principles of brain circuit function, investigating brain pathologies, and evaluating compounds for drug discovery.

## Introduction

The increase in patients with neurodegenerative and psychiatric disorders due to aging has made the development of therapeutic drugs for neurological diseases an urgent issue. As of 2023, it is estimated that approximately 55 million people worldwide have dementia (Hall et al., 2024), about 10 million have Parkinson’s disease (Marras et al., 2018), and around 280 million suffer from depression (Woody et al., 2017), making this a major social concern. Additionally, epilepsy is known to have a high incidence rate among the elderly (Sillanpää et al., 2011), and the number of epilepsy patients is expected to rise. One common brain function change observed in these neurological disorders is the abnormal release of neurotransmitters. Evaluating changes in neurotransmitter release serves as an important indicator for understanding disease mechanisms and for drug development.

Among the different types of dementia, Alzheimer’s disease, schizophrenia, and some forms of epileptic seizures are associated with the release of glutamate, one of the key neurotransmitters. The binding of glutamate to the N-methyl-D-aspartate (NMDA) receptor, a type of glutamate receptor, enhances excitatory transmission and plays a crucial role in higher brain functions such as memory and learning. However, in Alzheimer’s disease, excessive stimulation of NMDA receptors has been reported to induce cell death, leading to a decline in the number of neurons and resulting in memory and learning impairments. This phenomenon is known as glutamate excitotoxicity (Danysz and Parsons, 2012). In schizophrenia, a decrease in NMDA receptor function due to unknown causes leads to reduced activity of GABAergic neurons, triggering excessive firing of glutamatergic neurons and causing an overrelease of glutamate, which is believed to contribute to the onset of the disease (Nagai et al., 2017). Additionally, epileptic seizures are caused by abnormal synchronized firing of neurons. Normally, neurons maintain a balance between excitation and inhibition, but during seizures, this balance is disrupted. In particular, the reduced function of inhibitory neurotransmitters (such as GABA) leads to an increased release of excitatory glutamate (Wong, 2008). Measuring glutamate release is a valuable approach for understanding changes in neural activity, aiding in the elucidation of disease mechanisms and the development of therapeutic drugs for Alzheimer’s disease, schizophrenia, and epileptic seizures.

*In vitro* and *ex vivo* experiments offer several advantages, including the elimination of the need for large animal studies, high throughput, cost-effectiveness, and the ability to use disease-derived or healthy human cells. The evaluation of compounds using New Approach Methods (NAMs) has become an international trend (Bearth et al., 2025), and the demand for *in vitro* assessments is increasing in drug discovery, particularly in efficacy and safety testing (Liu et al., 2025). Methods for measuring glutamate *in vitro* include high-performance liquid chromatography (HPLC), fluorescence labeling assay, and electrochemistry measurement techniques. HPLC is a method that collects samples from cultured neuronal media or brain slice cultures and measures glutamate concentrations using chromatography (Zhang et al., 2005). Fluorescence labeling assay employ fluorescent glutamate indicators (such as iGluSnFR), allowing for the visualization of neurotransmitters with genetic and molecular specificity (Aggarwal et al., 2023). Electrochemistry measurement techniques utilize methods such as cyclic voltammetry (CV) and chronoamperometry (CA) to detect glutamate reduction currents (Dorozhko et al., 2015; Martínez-Perinán et al., 2023). For low-dose electrochemical detection of glutamate, enzymatic methods using glutamate oxidase can be employed, measuring reaction products such as hydrogen peroxide (H_2_O_2_) (Wang et al., 2024). Among these techniques, fluorescence labeling assays and enzyme-based electrochemistry measurements are the most suitable for real-time and non-invasive glutamate detection.

If real-time measurement of neurotransmitter release can be performed simultaneously with neural network activity, the relationship between neural activity changes and neurotransmitter release fluctuations can be clarified, leading to a deeper understanding of the underlying mechanisms and providing an effective evaluation method. One non-invasive, multi-site, high-temporal-resolution method for measuring *in vitro* neural network electrical activity is the microelectrode array (MEA) technique, which records field potential (FP) In our previous studies, we reported a technique for real-time electrochemistry measurement of dopamine using MEA chips (Suzuki et al., 2013). By using MEA to measure both FP and electrochemical reactions simultaneously, it becomes possible to non-invasively record both neural activity and neurotransmitter release. This approach eliminates the need to integrate expensive fluorescence microscopes required for fluorescence labeling assays, allowing for the measurement of two key indicators using a single device. In this study, we developed an MEA-based system capable of real-time simultaneous measurement of FP and electrochemistry (EC) signals. We designed an enzyme-modified carbon nanotube (CNT)-MEA chip capable of measuring both FP and electrochemical reactions of glutamate. Using the developed simultaneous FP/EC measurement system, we evaluated the glutamate sensitivity characteristics and conducted simultaneous measurements of FP and glutamate release from hippocampal brain slices.

## Materials and Methods

### Fabrication of Enzyme-Modified CNT-MEA

A microelectrode array (MEA; Alpha MED Scientific Inc., Japan) was used, consisting of 64 indium tin oxide (ITO) electrodes (size: 50 µm × 50 µm) arranged in an 8 × 8 square grid with a 150 µm spacing on a 5.0 cm × 5.0 cm glass substrate. Additionally, four reference electrodes (size: 200 µm × 200 µm) were separately positioned from the 64 recording electrodes, and CNT (Cup Stacked Carbon Nanotubes; GSI Creos, Japan) were electroplated onto the ITO electrode surface and reference electrodes. To prepare the CNT dispersion, CNT powder was dispersed in 1-methyl-2-pyrrolidone (NMP; Wako, Japan) using an ultrasonic homogenizer (Digital Sonifier 250; BRANSON, Japan). The prepared CNT dispersion was then mixed with sterilized water in a 1:1 ratio, and 2 mL of the mixture was dropped onto the electrode chip. Electroplating was performed using a DC stabilized power supply (ISO-TECH IPS303DD) by applying a voltage of 6.0 V for 60 seconds, repeated four times. Uniform electroplating was achieved by simultaneously applying voltage to all 64 electrodes using an MED connector (Alpha MED Scientific Inc.). For glutamate measurement, the CNT-MEA electrodes were modified with glutamate oxidase (Glu-Ox; Yamasa Corporation, Japan) and osmium-horseradish peroxidase (Os-HRP; BAS Inc., Japan) using glutaraldehyde as a crosslinker. 1% glutaraldehyde solution (pH 5.2, diluted with BSA) was mixed with Glu-Ox and Os-HRP in a 1:1:1 ratio, and 2 µL of the mixture was pipetted onto the electrode surface. The enzyme-modified electrodes were then stored in a light-shielded environment at 4°C overnight for drying. Before measurements, the electrodes were washed once with the measurement solution to remove excess reagents.

### Acquisition of SEM Images

The surface characteristics of electroplated CNTs were evaluated using a scanning electron microscope (SEM, SU8000, Hitachi, Japan). SEM images were acquired at an acceleration voltage of 3.0 kV and a magnification of 30,000 times.

### Simultaneous Measurement System for Field Potential (FP) and Electrochemistry (EC) Signals

The MEA64 system (MEA; Alpha MED Scientific Inc., Japan), which enables FP measurement, was customized by adding a circuit for chronoamperometry (CA) response measurement. In this system, one selected electrode among the 64 electrodes was used for CA measurement, while the remaining 63 electrodes were used for FP measurement. The CA measurement data and the FP data from one of the 63 selected electrodes were input into the multifunction I/O device (USB X SERIES Multifunction DAQ; National Instruments cpoe.) and the MED64 system. The FP data from the remaining 63 electrodes were directly input into the MED64 system. The working electrode was an arbitrarily selected enzyme-modified CNT microelectrode, the reference electrode was a set of four reference electrodes (size: 200 µm × 200 µm) plated with CNT electrodes, and the counter electrode was a platinum wire inserted into the solution. The sampling rate was set at 20 kHz, allowing for real-time simultaneous measurement of FP and glutamate reaction currents.

## Reagents

L-Glutamic acid (CAS RN: 56-86-0), hydrogen peroxide (H_2_O_2_, CAS RN: 7722-84-1), 25% glutaraldehyde solution (CAS RN: 111-30-8), potassium hexacyanoferrate (II) trihydrate, albumin from bovine serum (BSA, CAS RN: 13746-66-2), 1-methyl-2-pyrrolidone (NMP, CAS RN: 872-50-4), uric acid (CAS RN: 69-93-2), and caffeine (CAS RN: 58-08-2) were purchased from Wako pure chemical ocrporation (Japan). L-glutamate oxidase (Glu-Ox) was purchased from Yamasa Corporation (Japan). Osmium polymer/horseradish peroxidase (Os-HRP) was purchased from BAS Corporation (Japan).

Phosphate buffered saline tablets (PBS) were purchased from Takara bio inc (Japan). Dimethyl sulfoxide (DMSO, D2650), γ-aminobutyric acid (GABA, A2129), 4-aminoporidine (4-AP, 275875) and dopamine hydrochloride (DA, H8502) were purchased from Sigma. Platinum wire was purchased from Nilaco Corporation.

### Cyclic Voltammetry (CV) Measurement

To evaluate the sensitivity of CNT-plated microelectrodes, cyclic voltammetry (CV) measurements were performed using an HZ-7000 (Hokuto Denko, Japan). The measurement solution consisted of 2 mL of PBS and 10 mM potassium hexacyanoferrate (II) dissolved in PBS. CV measurements were conducted within a potential range of -1.0 V to 1.0 V (vs. Ag/AgCl reference electrode). The potential scan rate was set at 100 mV/s, with a sampling interval of 100 ms. All experiments were performed at room temperature.

### Chronoamperometry (CA) Measurement

Using the simultaneous FP/EC measurement system and HZ-7000, the CA response of the enzyme-modified CNT-MEA to glutamate and H_2_O_2_ was evaluated. PBS or artificial cerebrospinal fluid (aCSF) containing 124 mM NaCl, 3.0 mM KCl, 26 mM NaHCO_3_, 1.25 mM KH_2_PO_4_, 2.4 mM CaCl_2_/2H_2_O, 1 mM MgSO_4_/7H_2_O, and 10 mM D (+)-Glucose was used as the solvent. Glutamic acid solution or H_2_O_2_ solution dissolved in PBS or aCSF was administered cumulatively, and CA was measured at 0 V. In the simultaneous FP/EC measurement system, the enzyme-modified NT-MEA was used as the working electrode, the CNT electrode (size: 200 µm × 200 µm) was used as the reference electrode, and a platinum wire was used as the counter electrode, and measurements were made at a recording frequency of 20 kHz. In the HZ-7000, the enzyme-modified CNT-MEA was used as the working electrode, the Ag/AgCl wire was used as the reference electrode, and a platinum wire was used as the counter electrode, and measurements were made at a recording frequency of 10 Hz. All experiments were performed at room temperature. To demonstrate that the CA measurement results in mouse hippocampal slices are a response to glutamate, we confirmed the lack of reactivity with other neurotransmitters and inhibitors (GABA 10 μM, dopamine (DA) 100 nM, uric acid 10 μM, DMSO 0.1%, caffeine 200 μM). The experiment was a cumulative administration test performed using a simultaneous FP/EC measurement system.

### Reagent Simultaneous Measurement of FP and Glutamate in Acute Brain Slices

Acute brain slice experiments were conducted using 6- to 7-week-old male mice (C57BL/6NCrSlc, Japan SLC, Inc.). The mice were decapitated under isoflurane (Viatris Inc.) inhalation anesthesia, and the brain was extracted. The cerebellum and the anterior 1/3 to 1/4 of the cerebrum were removed. The brain was placed with the rostral side facing down, and approximately 20–30 degrees of the left and right hemispheres were trimmed. The trimmed brain was divided into right and left hemispheres, positioned dorsal side up, and sliced into 300 µm-thick sections using a slicer (Neo-LinearSlicer NLS-MT, DOSAKA EM CO., LTD.). The experiments were conducted following the animal experiment guidelines approved by Tohoku Institute of Technology (Approval Number: 2020-01). The brain slices were incubated for 1 hour at 30–32°C for recovery and maintained at room temperature until measurement. The recovery solution consisted of aCSF bubbled with a 95% O_ / 5% CO_ gas mixture. FP and electrochemistry measurements in mouse hippocampal slices were performed using the simultaneous FP/EC measurement system, with aCSF continuously perfused over the electrode surface while being bubbled with 95% O_ / 5% CO_ gas mixture. The electrode chip was placed on a heater (MED temperature-controlled connector MED-CP04H; Alpha MED Scientific Inc., Japan) set to 32°C. To observe changes in glutamate release, 200 µM caffeine was administered, and the responses were recorded.

### Analysis

For electrochemistry measurement data obtained using the HZ-7000, the calibration curve was generated by plotting the minimum currents recorded 20 seconds after the dropwise addition of glutamate or H_2_O_2_, over a 50-second interval. For electrochemistry measurement data obtained using the MED64 system connected to a multifunction I/O device, a 10-sample moving average was applied for smoothing, followed by downsampling from 20 kHz to 2 kHz. The calibration curve was calculated using the minimum currents recorded between 5 seconds and 47 seconds after glutamate administration. For electrochemistry measurement data obtained from simultaneous recordings in brain slices, the data was also downsampled from 20 kHz to 2 kHz. A fitting curve was applied within the selected data range to correct for slope variations. MATLAB and Excel were used for data analysis.

## Results

### Fabrication of Enzyme-Modified CNT-MEA

The electrode surface of the MEA used for FP measurements is typically coated with Pt/Pt-black to reduce impedance. However, for high-sensitivity detection of neurotransmitters via electrochemistry measurements, CNT was electroplated onto the ITO electrode surface. Figure 1A shows the entire view of the 64-electrode MEA probe along with phase-contrast images before and after electroplating (Fig. 1A). Scanning electron microscope (SEM) observations of the electrode surface after electroplating revealed that the CNT retained their tubular structure upon deposition (Fig. 1B). The increase in surface area resulted in enhanced electrochemical reaction sensitivity and reduced impedance, enabling the fabrication of an optimized electrode. To compare the electrochemical response of electrodes with and without electroplating, cyclic voltammetry (CV) measurements were performed using 10 mM potassium hexacyanoferrate (II). The CNT-electroplated electrode exhibited a significant reduction current compared to the bare ITO electrode, displaying a characteristic sigmoidal CV curve typical of microelectrodes. The peak current recorded at 0.337 V was 36.9 nA, with a current density of 1.48 nA/cm², confirming its high-sensitivity electrochemical properties (Fig. 1C).

**Figure 1.**
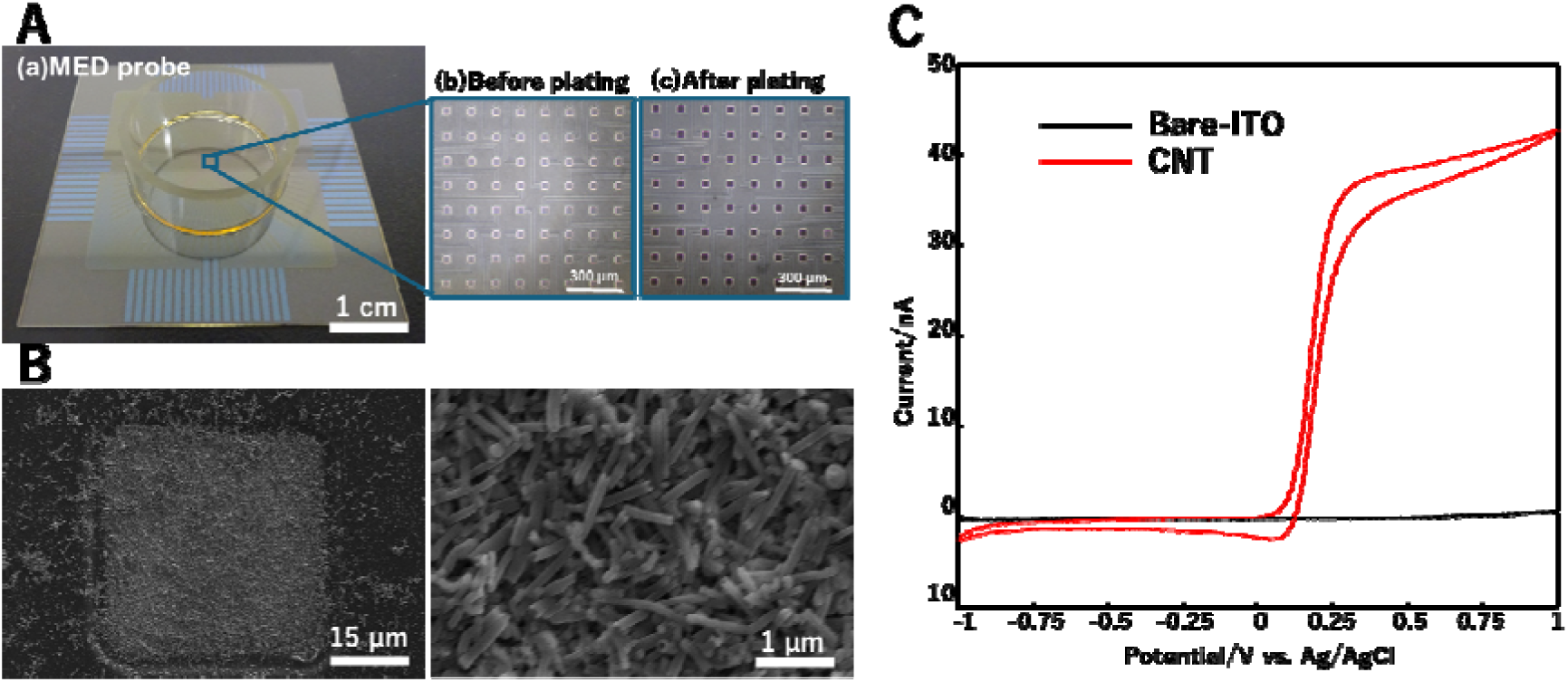
Electroplated CNT-MEA Chip and Its Electrochemical Properties. (A)(a) Overall view of the 64-electrode MEA probe. Scale bar = 1 cm. (b) Phase-contrast image of the 64 electrodes before CNT electroplating. (c) Phase-contrast image after CNT electroplating. Scale bar = 300 µm. (B) Scanning electron microscope (SEM) images of the CNT-microelectrode surface. (a) Scale bar = 15 µm. (b) Scale bar = 1 µm. (C) Cyclic voltammetry (CV) measurement results in 10 mM potassium hexacyanoferrate (II) (black: before electroplating, red: CNT-MEA).

### Development of a Simultaneous Measurement MEA System for Field Potential (FP) and Electrochemistry (EC) Signals

A measurement system was developed to enable the simultaneous recording of FP and EC using a CNT-MEA chip (Fig. 2A(a)). The CNT-MEA chip was placed on a heater (MED temperature control pad MED-CPB02; Alpha MED Scientific Inc., Japan), and signals were transmitted to the MEA64 system via the MEA connector (MAD-C03; Alpha MED Scientific Inc., Japan) before being output to a PC. The counter electrode was a platinum wire immersed in the liquid within the MEA probe. Figure 2A(b) illustrates the setup for simultaneous measurement of FP and EC. The MED64 system, which is originally designed for FP measurements, was customized by adding a circuit for electrochemistry measurements. In this setup one electrode out of the 64 electrodes was selected for electrochemistry measurement, while the remaining 63 electrodes were used for FP recording. The electrochemical currents from the measurement electrode and the FP data from a nearby selected electrode were input into a multifunction I/O device. The FP data from the remaining 63 electrodes were directly input into the MED64 system. This simultaneous FP/EC measurement system enabled real-time recording of both FP and EC. Additionally, to allow the CNT-MEA surface to detect glutamate, it was modified with enzymes (Glu-Ox, Os-HRP) and glutaraldehyde as a crosslinker (Fig. 2B(a)). The principle of glutamate measurement is based on the enzymatic reaction, where glutamate oxidase (Glu-Ox) converts glutamate into α-ketoglutarate, producing H_2_O_2_ as a byproduct. The generated H_2_O_2_ reacts with osmium-horseradish peroxidase (Os-HRP), and the resulting electrochemical current is measured (Fig. 2B(b)). To verify the functionality of the developed simultaneous FP/EC measurement system and its ability to simultaneously measure signals from biological samples, 100 nM glutamate was added every 100 seconds, and changes in FP recordings and chronoamperometry (CA) currents in EC recordings were evaluated. In the EC recording electrode, a reduction current was observed after each 100 nM glutamate addition. In contrast, the FP recording electrode only exhibited a liquid addition artifact, with no significant potential changes. These results confirmed that the enzyme-modified CNT electrode responds specifically to glutamate, and that the electrochemical reaction does not interfere with FP measurements (Fig. 2C).

**Figure 2.**
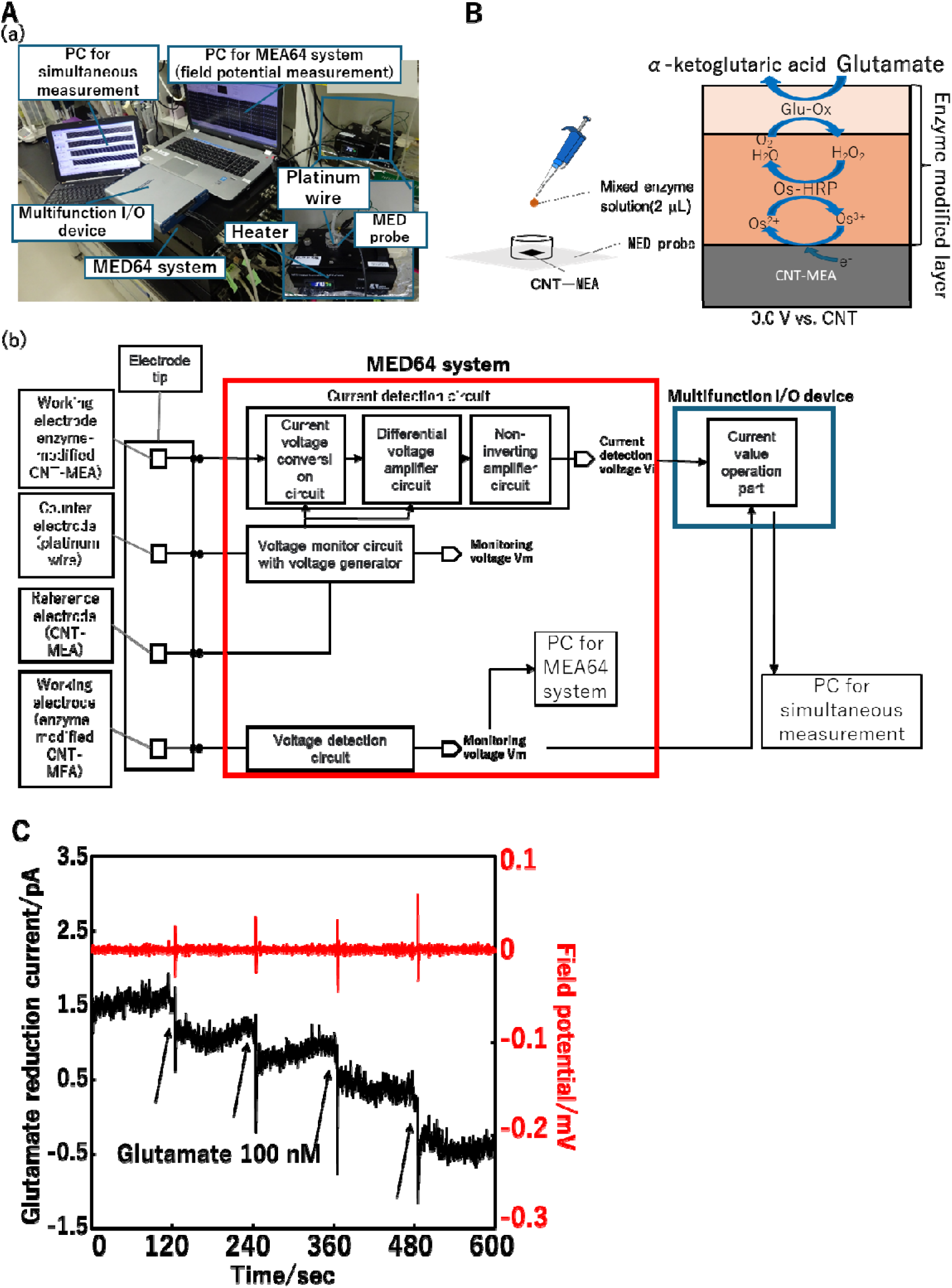
Simultaneous field potential (FP) and electrochemistry (EC) Measurement System and Glutamate Response. (A)(a) Experimental setup of the simultaneous FP/EC measurement system. (b) Circuit diagram of the simultaneous FP/EC measurement system. Red and blue square represent the MED64 system circuit and the multifunction I/O device circuit, respectively. (B) Conceptual diagram of the fabrication of the enzyme-modified CNT-MEA chip (left). Illustration of the glutamate reaction mechanism on the enzyme-modified CNT electrode. (C)Simultaneous measurement of FP and chronoamperometry (CA) at 0V during cumulative administration of 100 nM L-glutamate. Glutamate was administered every 100 seconds. Red: FP Black: CA

### Electrochemical Response to H_2_O_2_ in Enzyme-Modified CNT-MEA

To confirm whether Os-HRP detects H_2_O_2_ in a dose-dependent manner, the dose dependency and detection limit for H_2_O_2_ were evaluated. 10 nM H_2_O_2_ was sequentially added to the MEA probe containing artificial cerebrospinal fluid (aCSF), and the resulting changes in currents were analyzed. The current response changed stepwise as the H_2_O_2_ concentration increased (Fig. 3A). The response to 10 nM H_2_O_2_ exhibited linearity with a sensitivity of 0.207 pA/nM (n = 3, R² = 0.996, Fig. 3B). Similarly, when 100 pM H_2_O_2_ was sequentially added, the current response also changed stepwise with increasing H_2_O_2_ concentration (Fig. 3C). The response to 100 pM H_2_O_2_ showed a sensitivity of 14.4 fA/nM (R^2^ = 0.994, Fig. 3D). Since H_2_O_2_ was detectable at the 10 nM level, the enzyme-modified CNT-MEA possesses the necessary H_2_O_2_ sensitivity to measure glutamate release.

**Figure 3.**
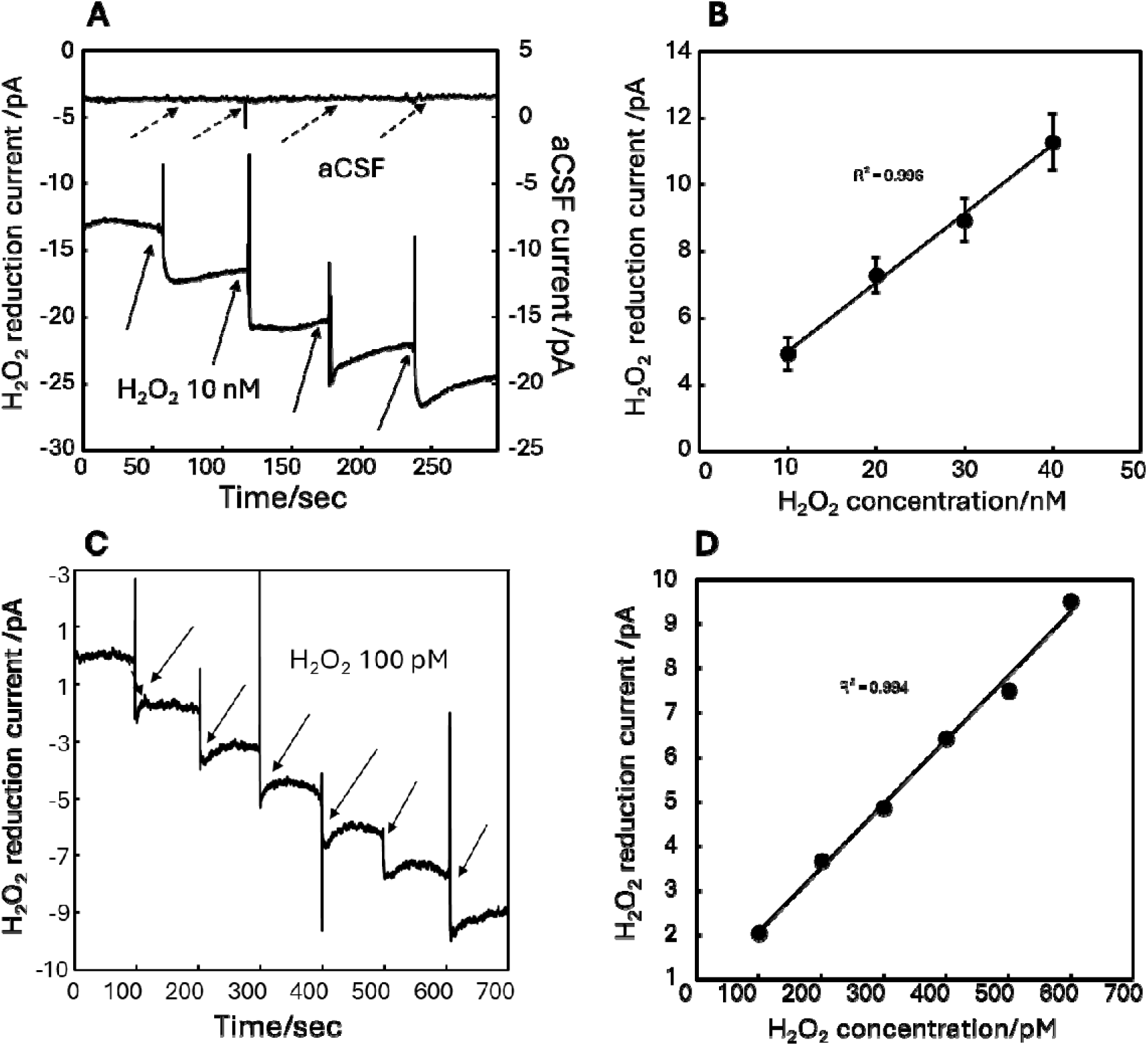
Electrochemical Response to H_2_O_2_ in Enzyme-Modified CNT-MEA Chip. (A) Changes in currents during cumulative administration of 10 nM H_2_O_2_. Arrows indicate the timing of administration. (B) Dose dependency of 10 nM H_2_O_2_ (n = 3). (C) Changes in currents during cumulative administration of 100 pM H_2_O_2_. (D) Dose dependency of 100 pM H_2_O_2_.

### Electrochemical Response to Glutamate in Enzyme-Modified CNT Electrodes

To evaluate the glutamate detection sensitivity of the fabricated enzyme-modified CNT electrodes, the dose dependency and detection limit were examined. Since the intraventricular concentration of glutamate ranges from 0.5 to 2 µM (Schultz et al., 2020), the dose dependency at the 100 nM level was measured. When glutamate was added every 100 seconds to the electrode probe containing aCSF, the current response changed stepwise (Fig. 4A). In contrast, no changes were observed when only the aCSF without glutamate was added. The response to 100 nM glutamate exhibited linearity with a sensitivity of 4.48 fA/nM (n = 3, R² = 0.999, Fig. 4B). Figure 4C show the current responses upon cumulative administration of 1 nM glutamate. The responses to 1 nM glutamate showed a sensitivity of 0.232 pA/nM (R^2^ = 0.955, Fig. 4D). Since glutamate release was detected in the range of a few nM to several hundred nM, it was confirmed that the electrode has sufficient sensitivity to detect glutamate in neuronal cell measurements.

**Figure 4.**
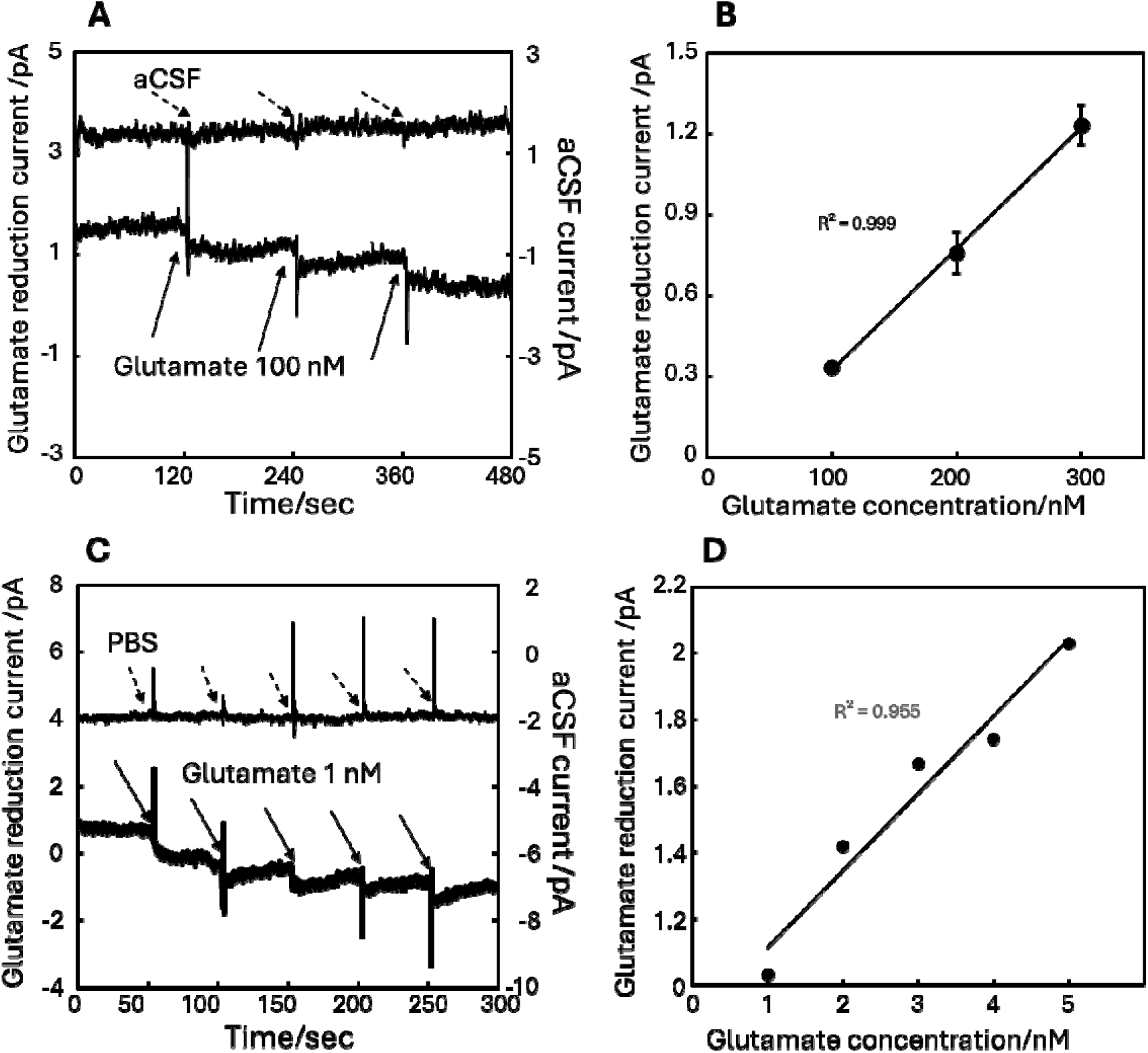
Electrochemical Response to Glutamate in Enzyme-Modified CNT-MEA Chip. (A) Changes in currents during cumulative administration of 100 nM glutamate. Arrows indicate the timing of administration. (B) Dose dependency of 100 nM glutamate (n = 3). (C) Changes in currents during cumulative administration of 1 nM glutamate. (D) Dose dependency of 1 nM glutamate.

### Electrochemical Response to Interfering Substances, Dopamine, and Glutamate

Since various substances are released from neural tissues, it was necessary to verify whether the currents obtained from the enzyme-modified electrode were specifically due to glutamate. Therefore, the effect of interfering substances was investigated. As shown in Figure 5, no changes in currents were observed when the aCSF or the neurotransmitter 10 µM GABA was added. Next, the administration of 100 nM dopamine resulted in an increase in currents by 4.50 pA, confirming that the electrode responded to dopamine independently of enzymatic activity. Additionally, since the response to dopamine was a positive oxidative current, distinct from the negative reductive current of glutamate, the two could be clearly differentiated. Furthermore, the administration of 10 µM uric acid, 0.1% DMSO, and 200 µM caffeine—all potential interfering substances released in the brain—caused no observable changes in currents. Finally, when 10 µM glutamate was administered, a reduction current was observed upon administration. These results indicate that the fabricated enzyme-modified CNT electrode can specifically detect both glutamate and dopamine while remaining unaffected by interfering substances.

**Figure 5.**
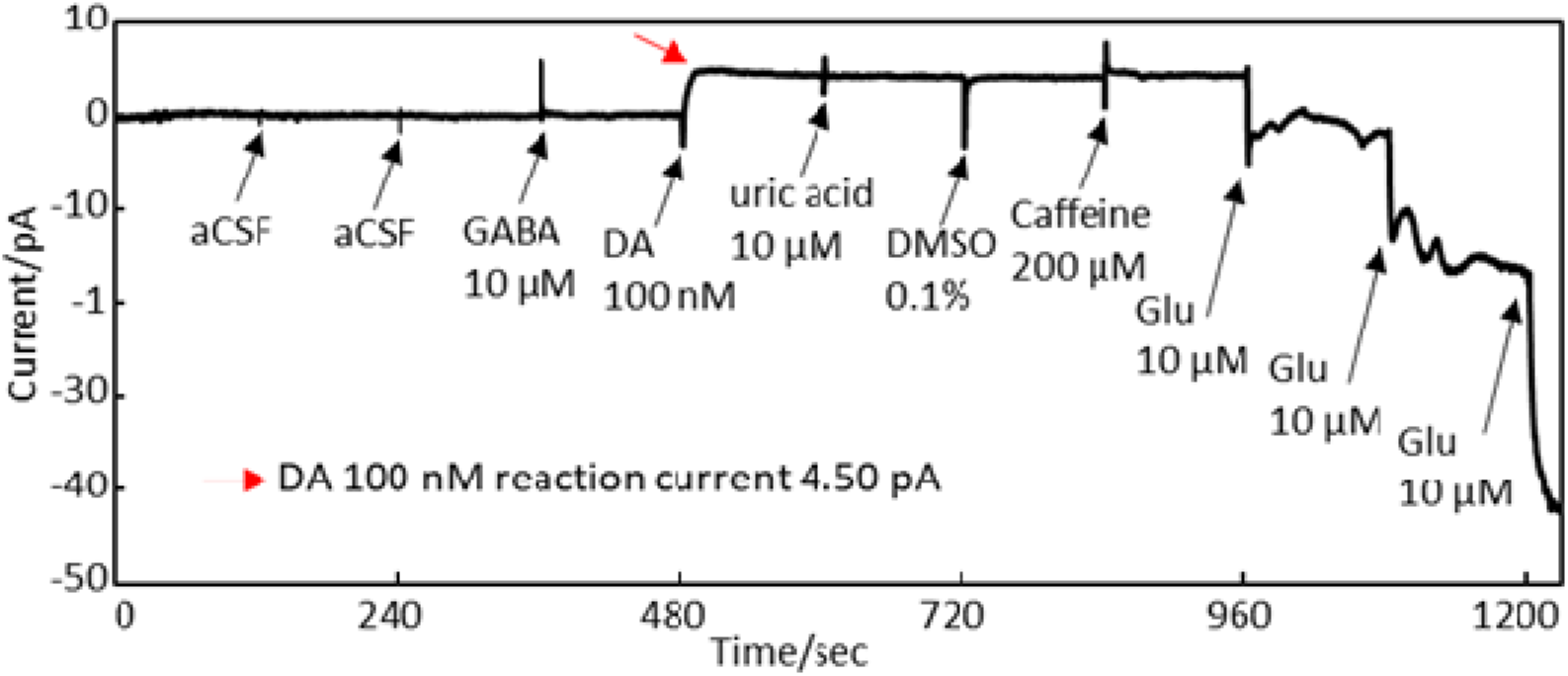
Evaluation of Responses to Interfering Substances, Dopamine, and Glutamate. CA measurements were performed during cumulative administration of aCSF, 10 μM GABA, 100 nM dopamine (DA), 10 μM uric acid, 0.1% DMSO, 200 μM caffeine, and 10µM glutamate.

### Simultaneous Measurement of Field Potential (FP) and Glutamate Release in Acute Brain Slices

Figure 6A shows a mouse hippocampal brain slice mounted on an enzyme-modified CNT-MEA chip. Chronoamperometry (blue square) was performed at the electrodes located between CA1 and CA2, and FP recording was performed at the remaining 63 electrodes. Figures 6B and 6C show the FP waveforms and the electrochemical waveforms of glutamate detection before and after caffeine administration. The FP waveform was obtained from the electrode (red square) located next to the CA measurement electrode. The oscillation activity of the hippocampal neural network, detected via FP, increased from 21 occurrences to 41 occurrences over 150 seconds after the administration of 200 µM caffeine (Fig. 6D). The spike-like signals in the currents (blue) in Figures 6B and 6C indicate that the neural activity detected in the FP is reflected in the currents. The glutamate responses were detected as downward slow waves. To quantify the reduction currents of glutamate, the oscillation-induced noise in the currents was smoothed and removed, and the area below 0 pA was calculated as the reaction charge. Before caffeine administration, glutamate release–induced current changes lasted 25.6 ± 10.2 seconds (n = 3). After caffeine administration, the current changes lasted 56.9 seconds.

**Figure 6.**
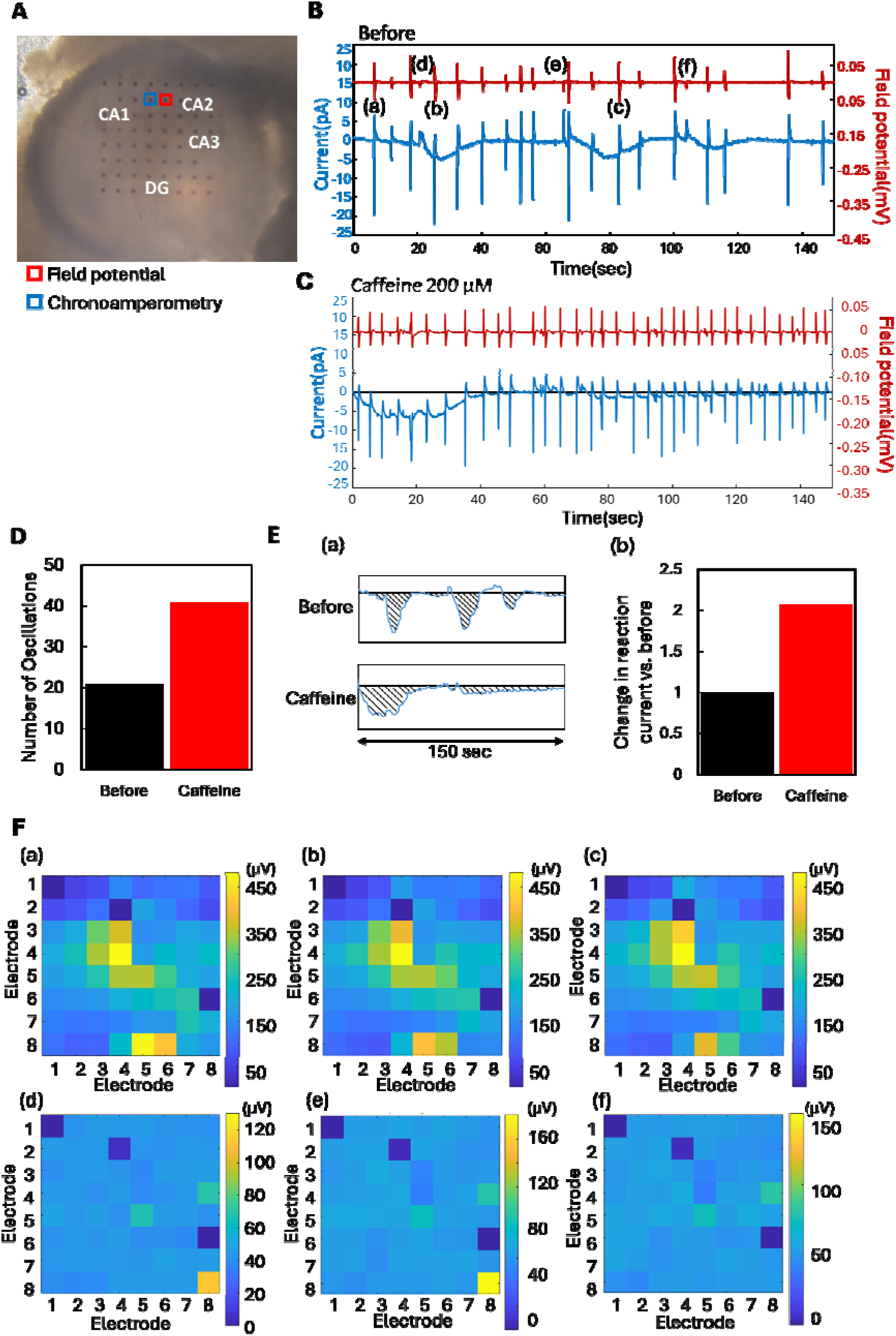
Simultaneous Measurement of Field Potential (FP) and Glutamate in Acute Brain Slices. (A) A mouse hippocampal brain slice mounted on an enzyme-modified CNT-MEA chip. The blue square shows a selected CA measurement electrode for glutamate reaction. The red squares indicate that FP waveforms were extracted from adjacent electrodes. (B) Simultaneous FP/EC measurement of FP (red) and glutamate reaction current (blue) during low-Mg aCSF perfusion. (C) Simultaneous FP/EC measurement of FP (red) and glutamate reaction current (blue) during the administration of 200 nM caffeine. (D) Comparison of oscillation counts in neural activities obtained from FP recordings. (E)(a) Conceptual diagram for the calculation of reaction charge for glutamate. (b) Comparison of glutamate reaction charge. (F) Heatmaps of the absolute values of field potential amplitudes mapped across 63 electrodes for oscillations (a)–(g).

The reaction charge over 150 seconds was 2.07 times higher after caffeine administration compared to before (Fig. 6E). These results indicate that changes in neural activity and neurotransmitter release before and after caffeine administration can be simultaneously measured using the FP/EC system. Even when FP oscillations were detected, no changes in glutamate-induced currents were observed in some cases. Based on these results, we hypothesized that the detection of glutamate release depends on differences in neuronal firing patterns. To test this hypothesis, we analyzed the oscillatory firing patterns (a) to (f) in Figure 6B using potential waveforms obtained from 63 electrodes. The absolute values of the potential waveforms were calculated, and heatmaps were generated for all 63 electrodes (Figure 6F). The heatmaps for oscillations (a) to (b), which did not exhibit current changes, were consistent across three trials. Similarly, the heatmaps for oscillations (d) to (f), in which current changes were observed, were also consistent across three trials. These findings suggest that whether glutamate-induced current changes are observed depends on differences in neuronal firing patterns.

### Simultaneous Measurement of Field Potential (FP) and Dopamine Release in Acute Brain Slices

Using the developed enzyme-modified CNT electrode, dopamine release from brain slices was successfully measured. Figure 7A shows the electrode location where dopamine was detected during 100 µM 4-aminopyridine (4-AP) administration and the hippocampal slice structure. In the CA2-CA3 region, nine upward oscillations were observed in the electrochemical measurement waveforms (blue, Fig. 7B) over 300 seconds, independent of the oscillation timing at 204 seconds detected in the FP waveform (red, Fig. 7B). For example, among the nine oscillations, the response occurring between 195.1 and 208.4 seconds lasted for 13.3 seconds. The peak current of oscillation at (a) in Fig. 7B was 19.8 pA. As shown in Figure 5, dopamine is detected as a positive current value. Additionally, the waveform recorded in this experiment was consistent with our previously reported dopamine release recordings from brain slices (Suzuki et al., 2013). Considering the recording location, it is likely that dopamine release originating from projections of the locus coeruleus was captured (Takeuchi et al., 2016). These results confirm that dopamine release from brain slices can be measured using the enzyme-modified CNT electrode.

**Figure 7.**
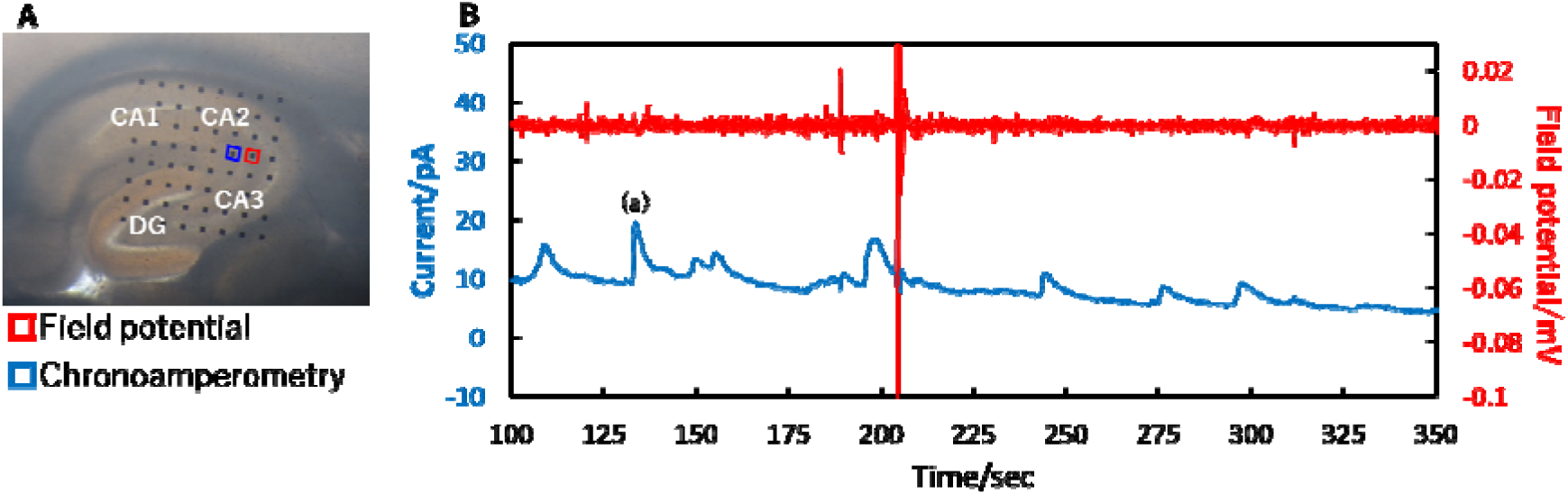
Simultaneous Measurement of Field Potential (FP) and Dopamine Release in Acute Brain Slices. (A) A mouse hippocampal brain slice mounted on an enzyme-modified CNT-MEA chip. The blue square shows a selected CA measurement electrode for dopamine reaction. The red squares indicate that FP waveforms were extracted from adjacent electrodes. (B) Simultaneous measurement of FP (red) and dopamine reaction current (blue) during 100 µM 4-AP administration in a brain slice.

## Discussion

We developed an MEA chip with electrochemical reactivity by electroplating CNT onto the MEA substrate. In the reaction of potassium hexacyanoferrate (II), a current density of 1.48 nA/cm² was recorded, demonstrating that the microelectrode exhibits high sensitivity to electrochemical reactions (Fig. 1). SEM images revealed that the CNT structure was maintained during electrodeposition, suggesting that the increased surface area contributes to enhanced sensitivity. Further improvements in sensitivity are expected through the selection of CNTs and optimization of electrodeposition conditions. A novel MEA system was developed using the CNT-MEA chip, enabling the simultaneous measurement of FP and EC signals. The simultaneous measurement of the electrochemical response to glutamate and FP demonstrated that FP could be stably recorded without being affected by the glutamate-induced current response (Fig. 2).

An enzyme-modified CNT-MEA was developed, demonstrating the capability to detect HLOL at concentrations of 100 pM and 10 nM, confirming its sufficient sensitivity for glutamate detection (Fig. 3). The difference in sensitivity between 10 nM and 100 pM (Fig. 3B and 3D) is considered to be influenced by the electrode fabrication conditions and enzyme activity. To achieve stable detection at the 100 pM level, further optimization of the CNT dispersion state, electroplating conditions, and enzyme coating conditions is necessary. Furthermore, the chip was confirmed to detect glutamate within a concentration range of a few nM to several hundred nM (Fig. 4). The detection limit for the glutamate response current was 1 nM, with a sensitivity of 0.232 pA/nM (Fig. 4D), which was approximately 52 times higher than the dose-dependent response at 100 nM (4.48 fA/nM, Figure 4B). Similar to H_O_ detection, further optimization of the CNT dispersion state, electroplating conditions, and enzyme coating conditions is expected to enable the fabrication of a highly sensitive electrode capable of stable glutamate detection at the 1 nM level.

The detection sensitivity of the developed glutamate sensor was compared with conventional glutamate sensors (Table 1). Reports of enzyme-modified electrodes using the same Glu-Ox enzyme for chronoamperometry (CA) measurements indicate a glutamate detection range of 0.01–1400 µM (Batra and Pundir, 2013; Chen et al., 2020; Hamdi et al., 2006; Hughes et al., 2016; Maity and Kumar, 2019; Mentana et al., 2020; Niwa et al., 1996; Razumiene et al., 2023; Scoggin et al., 2019; Wang et al., 2020). The highest reported detection sensitivity was achieved using a peptide aptamer-based sensor, with a linear detection range of 0.01–1 µM and a detection limit of 0.0072 µM (Niwa et al., 1996). The sensor developed in this study exhibited a linear detection range of 0.1–0.3 µM and a detection limit of 0.001 µM, which is comparable to the peptide aptamer-based sensor. However, glutamate measurement from cellular release has not been reported, and stability remains a challenge. Sensors using Glu-Ox-modified electrodes for cyclic voltammetry (CV) measurements have been reported with detection ranges of 0.5–10,000 µM (Deng et al., 2013) and 0.02–500 µM (Batra et al., 2016). However, CV measurements require varying the applied voltage, which may interfere with FP recordings during simultaneous measurements. Electrodes using L-glutamine-binding protein (QBP) as the enzyme operate within a relatively narrow scanning voltage range (-0.1 to -0.2 V) (Takamatsu et al., 2021). Additionally, some reports describe non-enzymatic sensors (Wang et al., 2022; Wang et al., 2024), but they also require CV measurements. Besides Glu-Ox, glutamate dehydrogenase has been frequently used as an enzyme in biosensors (Chakraborty and Raj, 2007; Doaga et al., 2009; Girousi et al., 2001; Hughes et al., 2015; Liang et al., 2015; Martínez-Perinán et al., 2023; Martinez-Perinan et al., 2016; Meng et al., 2009). The lowest glutamate concentration detected using a glutamate dehydrogenase-based sensor was 0.1 µM (Meng et al., 2009). However, Glu-Ox generally provides higher sensitivity across various studies (Table 1). The enzyme-modified CNT-MEA developed in this study exhibited a detection sensitivity in the range of a few nM to several hundred nM, which is sufficiently high for measuring glutamate in biological samples. Additionally, it allows measurement without applied voltage, making it ideal for simultaneous recording of cellular glutamate release and FP, which is a key advantage of combining CNT-MEA with enzyme modification.

**Table 1.**
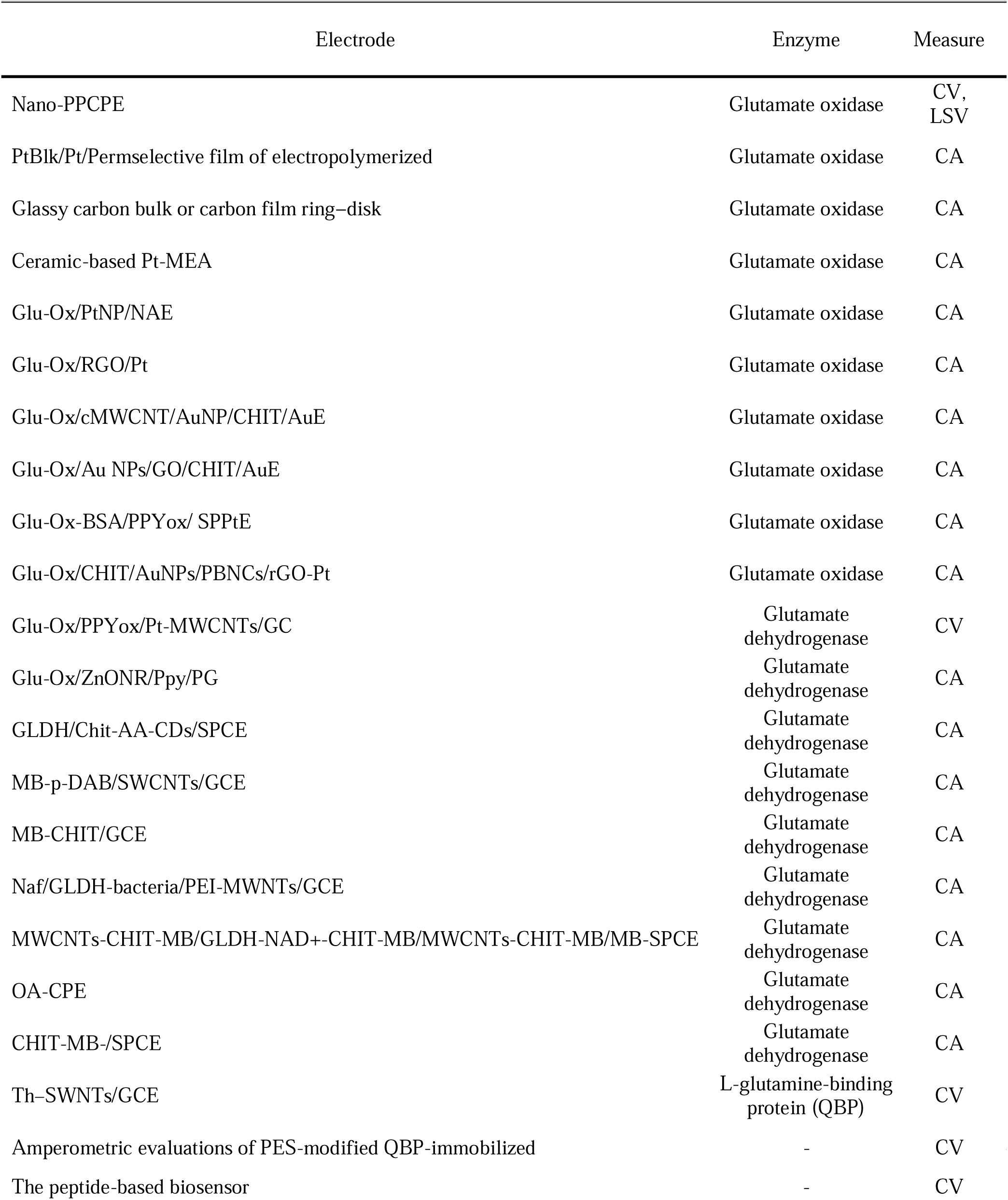
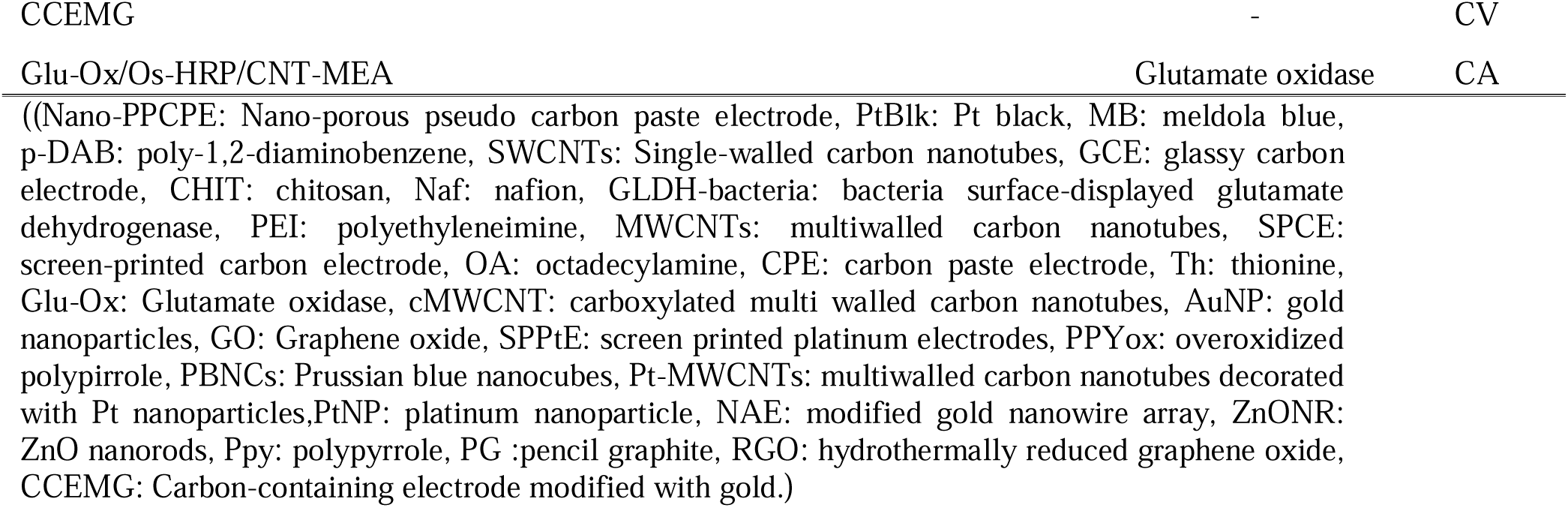
Comparison of Glutamate Biosensors.

Furthermore, using the developed enzyme-modified CNT-MEA and the simultaneous FP/EC measurement system, we successfully recorded FP and glutamate release from hippocampal brain slices and detected the effects of caffeine. In hippocampal brain slices, current changes associated with glutamate release were detected for approximately 20 seconds following oscillations (Fig. 6). A previous study targeting the dentate gyrus of the rat hippocampus reported that an enzyme-based electrochemical sensor detected glutamate releases lasting approximately 30 seconds following KCl stimulation (Hu et al., 1994). The current changes observed in our study were consistent with these findings, suggesting that the slow-wave phenomenon observed is characteristic of enzyme-based electrodes. Additionally, the similarity in waveform patterns between our study and the previous study further supports that the detected signal corresponds to glutamate release from brain slices.

Following caffeine administration, significant current changes were observed up to approximately 40 seconds, with the reaction charge increasing by a factor of two. This is likely due to an increase in neural activity, leading to an increased release of glutamate from synapses. A previous *in vivo* study using microdialysis in rats reported that systemic administration of caffeine increased extracellular glutamate concentration by approximately 50% in the medial shell of the nucleus accumbens (Solinas et al., 2002). Subsequent reductions in glutamate release were consistent with previous findings that glutamate release decreases through adenosine receptor-mediated mechanisms (Marcoli et al., 2003). The ability to measure glutamate in real-time using this technique allowed for the detection of temporal changes in glutamate release. Through simultaneous measurement with FP and CA, it was found that not all oscillations corresponded to glutamate responses (Fig. 6). As shown in the heatmaps in Fig. 6F, the difference in neural circuit firing patterns determined whether glutamate release could be detected. Since the brain slice experiments were conducted under perfusion conditions, it is also possible that the direction of perfusion influenced glutamate detection. Future studies will include electrical stimulation experiments to further investigate the relationship between firing patterns and glutamate release, as well as the impact of perfusion direction on detection accuracy.

Dopamine release from hippocampal brain slices was also recorded. The administration of 4-AP is presumed to have increased neural activity not only in the hippocampus but also in the surrounding tissues. The upward current signals in Fig. 7 are believed to reflect dopamine release from projections of the locus coeruleus. This assumption is based on the fact that the timing of hippocampal oscillations detected in extracellular recordings did not coincide with the timing of dopamine release (Fig. 7B). A 2016 study reported that dopaminergic projections from the locus coeruleus to the hippocampus play a role in memory consolidation (Takeuchi et al., 2016). Since the developed method enables simultaneous FP/EC measurement of long-term potentiation testing via FP recording, dopamine release, and glutamate release, it is expected to become an effective tool for elucidating the mechanisms of memory consolidation.

In this study, we developed an MEA system capable of simultaneously measuring FP and EC signals. Additionally, we developed an enzyme-modified CNT-MEA capable of detecting glutamate at concentrations ranging from several nM to several hundred nM. Using the developed enzyme-modified CNT-MEA and the simultaneous FP/EC measurement system, we successfully recorded field potential and glutamate release from hippocampal brain slices, detected the effects of caffeine, and demonstrated the ability to detect dopamine release. We believe that the findings of this study, which enable the simultaneous measurement of neural activity, glutamate release, and dopamine release, will contribute to a deeper understanding of brain circuit mechanisms, pathological brain conditions, and compound evaluation for drug discovery.

## Acknowledgments

The simultaneous FP/EC measurement equipment for this study was developed in collaboration with Hideyasu Jiko in Alpha MED Scientific Inc. and Shun Nakajima in SCREEN Holdings Co., Ltd. This study was supported by JKA Foundation Grant Numbers 24MC1007-136.

## CRediT authorship contribution statement

Aiko Hasegawa: Data curation, Formal analysis, Investigation, Methodology, Validation, Visualization, Writing – original draft, Writing – review & editing. Naoki Matsuda: Data curation, Formal analysis, Investigation, Methodology, Resources, Software, Visualization, Writing – original draft, Writing – review & editing. Ikuro Suzuki: Conceptualization, Data curation, Funding acquisition, Investigation, Methodology, Project administration, Resources, Supervision, Visualization, Writing – original draft, Writing – review & editing.

## Declaration of competing interest

The authors declare that they have no known competing financial interests or personal relationships that could have appeared to influence the work reported in this paper.

## Notes

### Competing Interest Statement

The authors have declared no competing interest.

